# Differential strengths of molecular determinants guide environment specific mutational fates

**DOI:** 10.1101/134569

**Authors:** Rohan Dandage, Rajesh Pandey, Gopal Jayaraj, Kausik Chakraborty

## Abstract

Under the influence of selection pressures imposed by natural environments, organisms maintain competitive fitness through underlying molecular evolution of individual genes across the genome. For molecular evolution, how multiple interdependent molecular constraints play a role in determination of fitness under different environmental conditions is largely unknown. Here, using Deep Mutational Scanning (DMS), we quantitated empirical fitness of ∼2000 single site mutants of Gentamicin-resistant gene (GmR). This enabled a systematic investigation of effects of different physical and chemical environments on the fitness landscape of the gene. Molecular constraints of the fitness landscapes seem to bear differential strengths in an environment dependent manner. Among them, conformity of the identified directionalities of the environmental selection pressures with known effects of the environments on protein folding proves that along with substrate binding, protein stability is the common strong constraint of the fitness landscape. Our study thus provides mechanistic insights into the molecular constraints that allow accessibility of mutational fates in environment dependent manner.

**Author Summary:** Environmental conditions play a central role in both organismal adaptations and underlying molecular evolution. Understanding of environmental effects on evolution of genotype is still lacking a depth of mechanistic insights needed to assist much needed ability to forecast mutational fates. Here, we address this issue by culminating high throughput mutational scanning using deep sequencing. This approach allowed comprehensive mechanistic investigation of environmental effects on molecular evolution. We monitored effects of various physical and chemical environments onto single site mutants of model antibiotic resistant gene. Alongside, to get mechanistic understanding, we identified multiple molecular constraints which contribute to various degrees in determining the resulting survivabilities of mutants. Across all tested environments, we find that along with substrate binding, protein stability stands out as the common strong constraints. Remarkable direct dependence of the environmental fitness effects on the type of environmental alteration of protein folding further proves that protein stability is the major constraint of the gene. So, our findings reveal that under the influence of environmental conditions, mutational fates are channeled by various degrees of strengths of underlying molecular constraints.

## Introduction

Continuously fluctuating natural environments are known to a play central role in natural selection by constituting environment to phenotype interactions (E2P). Such interactions can predispose a particular genotype to alternative fates through diverted evolutionary trajectories (1–5). At lower scales, molecular evolution of cellular genes, underlie the emergent organismal adaptations. Cellular responses to the environmental conditions are well known to be often enabled by maintenance of normal proteostasis (6). Therefore, an investigation of whether and to how much extent protein folding and stability play a role in guiding mutational fates would be a key factor in the efforts to elucidate molecular level E2P interactions.

Considering low rates of spontaneous mutations across variety of organisms (7), single site mutations provide a view of immediate next fates of a gene. DMS approach has enabled scanning of large scale of mutations in a high throughput manner (8,9); allowing comprehensive analysis of sequence-space of genes. Resultant Distributions of Fitness Effects (DFE) provide a continuous series of fitness effects ranging from strongly deleterious to beneficial which is a valuable resource for quantitative genetics (10). In recent years, exploration of environmental effects with a large-scale genotype to phenotype (G2P) data (11,12) has resulted in the identification of environment specific differential mutational sensitivity. However, qualitative and quantitative identification of determinants of fitness effects has been a challenging task (13). In addition to improve much needed mechanistic understanding (14) of E2P interactions, such information can potentially increase the robustness in current approaches of prediction of phenotypes from genotypic information(15,16).

On a fitness landscape of a gene, levels of fitness of mutations may depend on number of molecular constraints which would shape the landscape. Here, to systematically investigate the underlying mechanisms of E2P interactions, firstly, we monitored fitness landscapes of Gentamicin (Gm) resistant gene GmR (aminoglycoside 3-N-acetyltransferase (aacC1)) under different physical and chemical environmental conditions which can have differential fitness effects. Among physical environments, we studied effects of lower (30°C) and elevated (42°C) temperature than the normal growth temperature of *E. coli* i.e. 37°C. Elevated temperature condition is known to limit foldability of temperature-sensitive mutants (17), hence its influence on the fitness landscape of GmR allows us to understand the implication of proteostasis on limiting the sequence space for this protein. Among chemical environments, we studied two model chemical chaperones - TMAO and glycerol which are known to assist cellular protein folding in vitro and in vivo, assist mutational buffering (18) and thermodynamically act on early refolding intermediate by non-identical mechanisms (19).

In this study, we show that mutations may provide selective advantage or disadvantage depending on acting environmental conditions. Among multiple molecular constraints of fitness landscape of GmR, perturbation of stability is identified to be the universal strong constraint across all test environments. Except for the case of simultaneous action of multiple individual environments, the relative selection pressures conferred by physical and chemical environments are largely dependent on folding constraint and hence are predictable. For instance, elevated temperature imposes a negative selection pressure while chemical chaperones are found to exert mutational robustness (buffering effect). Collectively, through mutational scanning of an essential gene, this study uncovers the largely unclear role of environments in protein evolution and contributions of underlying molecular constraints in determining the mutational fates.

## Results

### Deep mutational scanning of GmR

We implemented DMS method (8) to monitor survivabilities and empirical competitive fitness levels of individual mutants from a Single Site Mutation (SSM) library of GmR (18). For co-culture bulk competitions, we used purifying selection i.e. Gm concentration of 12.5 μg/mL which is ∼4 fold lower than the minimal inhibitory concentration (MIC) of *E. coli* (K-12) expressing wild type GmR while being higher than the MIC of the host system i.e. *E. coli* alone (S1 Fig 1A). While survivability of the antibiotic resistance is dosage dependent (S1 Fig 1B), use of a weaker Gm selection level allows assessment of mutants with moderate catalytic advantage which would otherwise be lost because of higher likelihood of ‘quick fix’ outcomes (20). Note that when not otherwise mentioned, for the bulk competitions, 12.5 μg/mL Gm is used. Co-culture bulk competitions of mutants were carried out in two sets. One in presence of Gm selection (selected pool) and other in absence of Gm selection (unselected pool). At the end of the bulk competitions, ultra-deep sequencing of the amplicons of GmR allowed quantitation of counts of mutants survived at the end of each competition (Fig 1A). To understand E2P effects of different environments, co-culture bulk competitions were carried out under different test environmental conditions (Materials and Methods). 37°C being the optimal growth temperature of *E. coli*, we simply refer it as the reference environment. When not otherwise noted, test environments would be compared against the reference environment (37°C, 12.5 μg/mL Gm).

**Fig 1.**
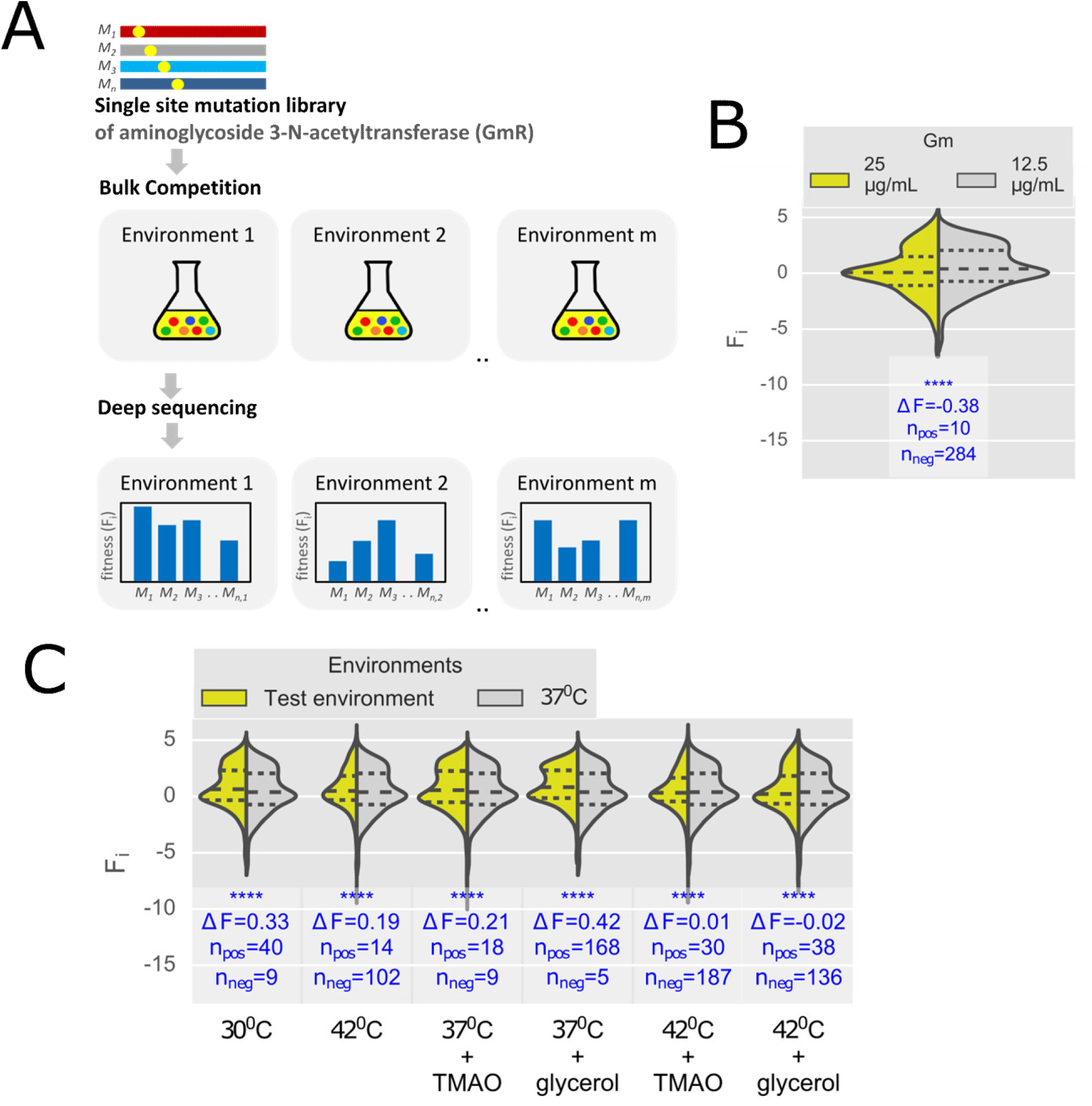
Deep mutational scanning of GmR. (A) Experimental strategy to monitor survivabilities and competitive fitness of GmR mutants (Materials and Methods). SSM library of GmR was subjected to co-culture bulk competition in different environments in presence of Gm selection (selected pool). An independent co-culture bulk competition in the absence of Gm selection (unselected pool) acts as a reference condition. Preferential enrichments in counts of mutants - measured via deep sequencing - in selected with respect to that unselected pool provides a measures of the survivabilities and proxies for competitive fitness levels of mutants. (B) DFE obtained at stringent concentration of Gm i.e. 25μg/mL (37°C) is compared to DFE obtained at 12.5 μg/mL (37°C). (C) Similarly, DFEs obtained under physical and chemical test environments and combinations of environments are compared to reference environmental condition (37°C). Here, all the bulk competitions were carried out at purifying selection of 12.5 μg/mL Gm. F_i_ denotes fitness score. Histograms of DFEs are fitted by kernel density estimation. Heights of the DFEs are in scale with the frequencies of mutants. Dashed lines represent quartile values of the fitness per distribution. Level of significance is measured by two sided Wilcoxon signed-rank test. Relative selection pressure imposed by a test environment is with respect to control condition is measured in terms of difference between average fitness scores (ΔF) and number of mutants with higher (n_pos_) or lower (n_neg_) survivability than control environment.

Strong correlations between counts of the mutants from independent biological replicates of bulk competitions (S1 Fig 2) indicate low level of biological noise associated with experiments. Preferential enrichments i.e. log fold changes of the counts of the mutants in selected pool with respect to that in unselected pool serve as proxies for competitive fitness of mutants. Next, we implement a strategy adopted in previous study on similar mutational scanning of antibiotic resistant gene (21), to classify statistically neutral mutations by applying a cutoff of μ±2σ (μ: mean, σ: standard deviation) obtained from fitness of unselected pool (S1 Fig 3). Accordingly, mutants were classified as relatively enriched (*F*_*i*_>μ+2σ) and depleted (*F*_*i*_<μ-2σ). Fitness levels of synonymous mutations across different environments are found to strongly correlate (Pearson’s *r* > 0.88, S1 Fig 4) with that fitness of synonymous mutations under reference environment (37°C, 12.5 μg/mL Gm); thus diminishing the influence of codon bias associated with G2P interactions of GmR. Therefore, unless specified, in the subsequent analysis of mutational data, we primarily utilize non-synonymous mutants. This way, from a co-culture bulk competition under reference conditions (37°C, 12.5 μg/mL Gm), we obtained empirical fitness of 2004 non-synonymous mutants (n) (S1 Fig 5, S2 Table). Among them, 609 mutants are enriched, 262 mutants are depleted and 1133 are classified as statistically neutral (S1 Table1).

**Fig 2.**
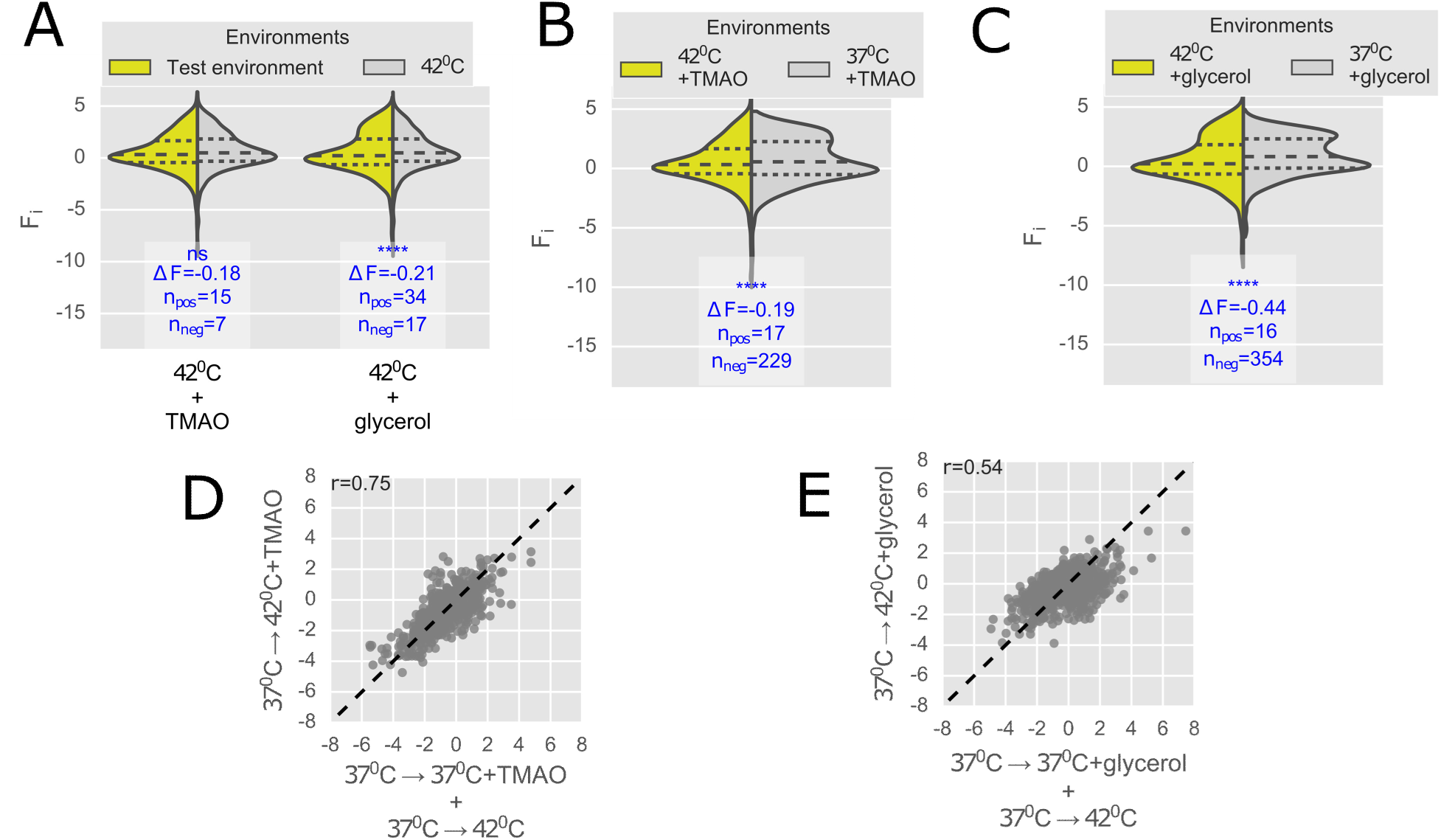
Selection strengths of combination of environments with respect to individual constituent environments. (A) In order to determine the relative selection strength of constituent elevated temperature condition in the combinations, DFEs of combination of environments i.e. 42°C+TMAO and 42°C+glycerol are compared against the DFE at 42°C. Similarly, DFE obtained under 42°C+TMAO condition is compared against TMAO condition (B) and DFE under 42°C+ glycerol condition is compared against glycerol condition (C). *F*_*i*_ denotes fitness score. Histograms of DFEs are fitted by kernel density estimation. Heights of the DFEs are in scale with the frequencies of mutants. Dashed lines represent quartile values of the fitness per distribution. Level of significance is measured by two sided Wilcoxon signed-rank test. Relative selection pressure imposed by a test environment is with respect to control condition is measured in terms of difference between average fitness scores (ΔF) and number of mutants with higher (n_pos_) or lower (n_neg_) survivability than control environment. Correlations between change in fitness of mutants under combination of environments i.e. (D) 42°C+TMAO and (E) 42°C+glycerol relative to reference environment with addition of fitness changes for constituent environments relative to reference environment. Dashed line denotes equal values of x and y variables. r is Pearson’s correlation coefficient.

**Fig 3.**
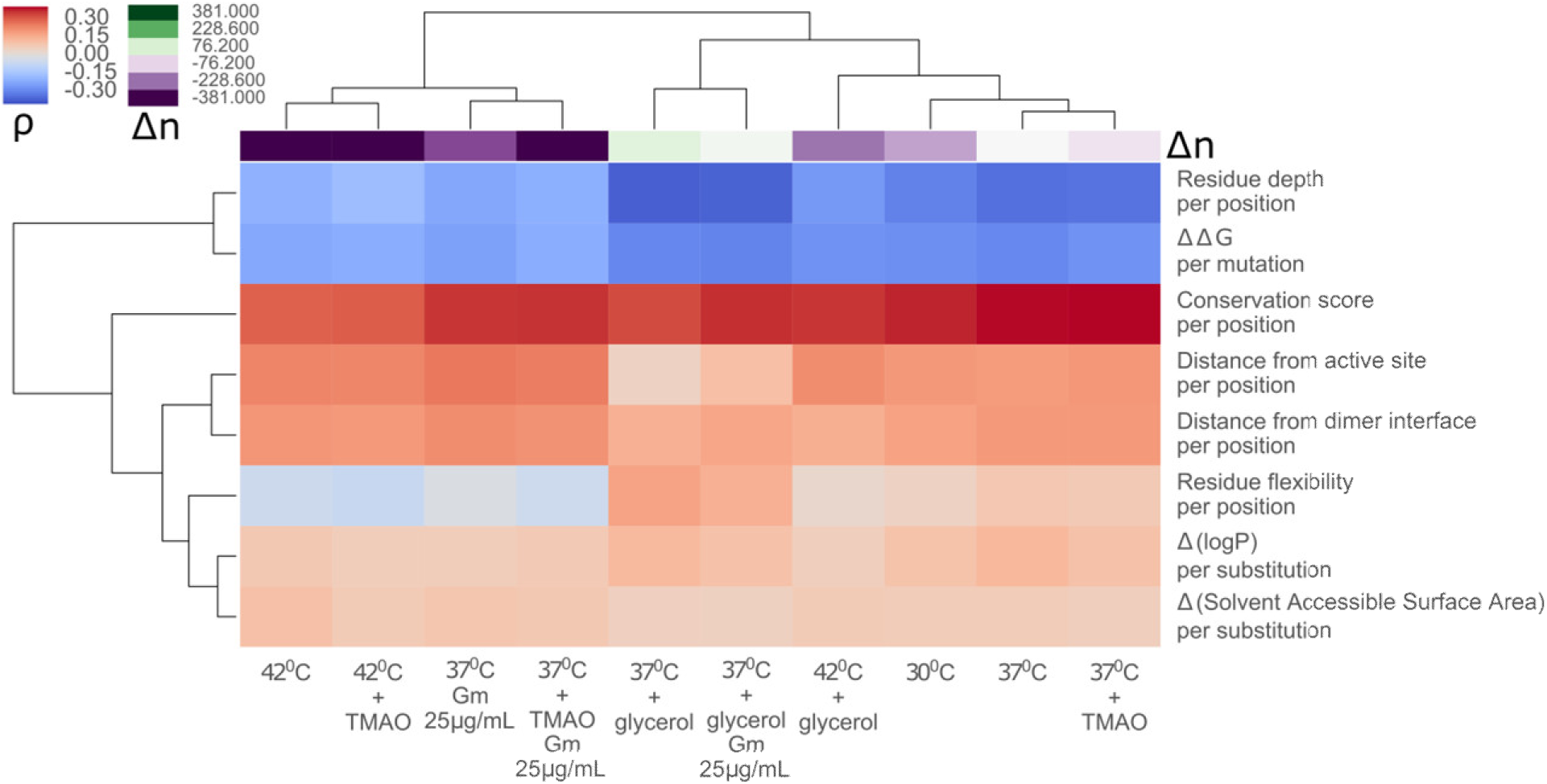
Differential strengths of molecular constraints underlying environmental fitness effects. Fitness scores obtained under different environmental conditions are correlated with a set of molecular features. Each locus in the heatmap indicates Spearman’s rank correlation coefficient between fitness scores of an environment (in columns) and a molecular feature (in rows). ρ is Spearman’s rank correlation coefficient and Δn is difference between number of mutants survived in test environments with respect to reference environment (37°C). Euclidean clustering along rows and columns is based on the Spearman’s rank correlation coefficients.

**Fig 4:**
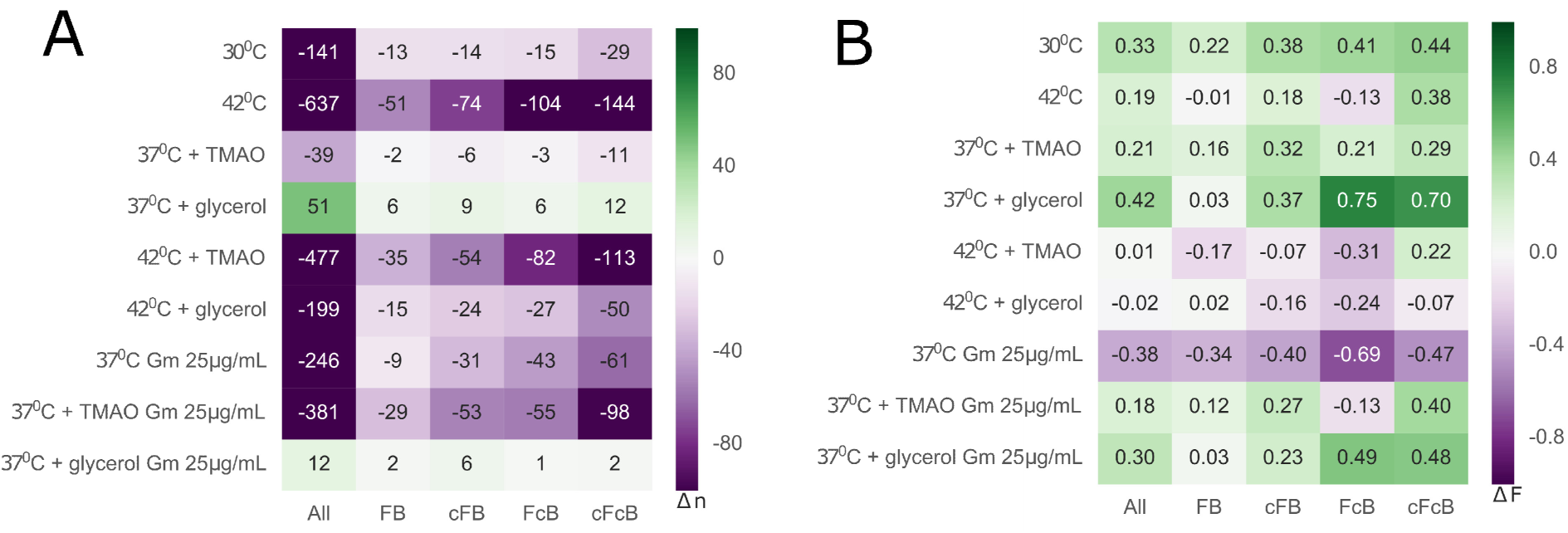
Survivability and fitness of mutants show dependence on perturbation of folding and binding. Subset-wise relative selections imposed by test environments with respect to reference environment i.e. 37°C at 12.5μg/mL Gm is measured in terms of (A) difference in number of survived mutants (Δn) and (B) difference in average fitness of mutants (ΔF) with respect to that of reference environment. ‘All’ denotes all mutations in a given the environmental condition. Among subsets of mutants (FB, cFB, FcB and cFcB), F and B represent mutants with low ΔΔG and high distance from active site respectively, whereas cF and cB represent mutants with high ΔΔG and low distance from active site respectively.

**Fig 5.**
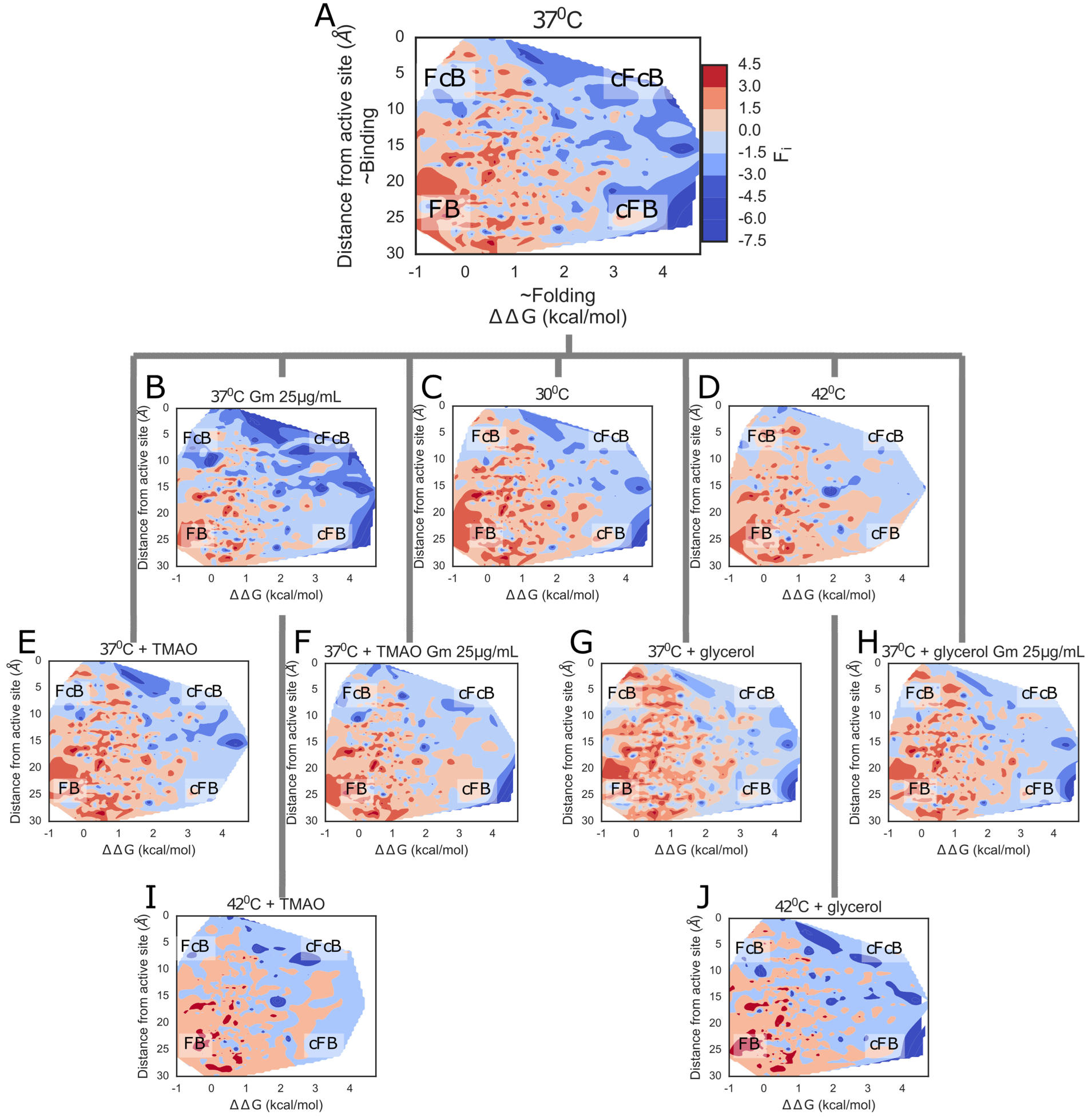
Unique reshapings of environment specific fitness landscapes. Environment specific fitness effects are visualized in a form of a fitness landscapes (A-J) lined by folding (ΔΔG) and binding (distance from active site) components with Z-component of the plot scaled by fitness levels (F_i_). Contour surfaces are generated by nearest neighbor interpolation. Regimes at the corners of the fitness landscapes thus represent subsets of mutants based on folding and binding components i.e. FB, cFB, FcB and cFcB. Colors of all contour plots are scaled according to the colorbar associated with panel A.

We compared pairs of DFEs obtained for various test environments by a metric defined as a difference in average fitness scores of mutants between test and reference condition (ΔF) which indicates the relative increase or decrease in fitness of surviving mutants in test condition with respect to reference environment. Additionally, number of mutants having significantly higher or lower fitness in test environment are measured (n_pos_ and n_neg_ respectively). Here, to account for the inherent experimental and biological noise, a cutoff is assigned for defining significant increase or decrease in the fitness over the inherent dispersion within replicates (Materials and Methods).

DFE of GmR obtained at 25 μg/mL Gm (37°C) reveals a depletion of fitness with a skew towards the deleterious fitness levels as compared to DFE obtained at 12.5 μg/mL Gm (Fig 1B). The lowered average fitness (ΔF = −0.38) is accompanied by lowered survival of mutants in this condition (Δn = −246). Such dosage dependent deleterious fitness effects are expected from increased stringency for *in vivo* catalytic functionality and survivability of mutants which is consistent with previous reports on mutational scannings of other antibiotic resistant genes (21–23). In terms of distribution of enriched and depleted mutants per position, mutations at sites with low evolutionary rate, such as binding sites, show depleted levels of fitness while mutations at sites with high evolutionary rate, specifically at N-terminal region, exhibit high fitness levels (S1 Fig 6). Collectively, dosage dependence and consistent trends with conservation scores support that the observed empirical fitness values are representatives of the *in vivo* functionalities of GmR mutants. This further allows us to ascribe empirically quantified fitness values to the *in vivo* functionalities of the mutants.

### Environmental conditions induce variable fitness effects

Using our experimental system, to understand E2P interactions, we monitored effects of sets of physical and chemical environments. In our study, low (30°C) and elevated (42°C) temperature conditions comprising a set of physical environments, while treatments of chemical chaperones TMAO and glycerol constitute a set of chemical environments. Physical environments especially elevated temperatures are well known to have limiting influence on the mutational tolerance (17). On the other hand, in case of chemical environments, solvent accessible surface area is known to be one of the strong predictor of protein evolution rate (24–26), suggesting that solvent-protein interactions are critical for the understanding of molecular evolution. In our study, we monitor effects of TMAO and glycerol which are known to act as chemical chaperones in assisting folding of proteins by non-identical mechanisms (19) as well as assisting buffering of genetic mutations (18).

We utilize prior information gained from growth kinetics of the model system i.e. wild type GmR harboring *E. coli* under different test environments to establish the strengths of selection pressures relative to reference environment. As compared to normal growth temperature of *E. coli* (37°C), both lower (30°C) and elevated temperatures (42°C) show non-optimal growth (shown in S1 Fig 7A). At elevated temperature, the amplitude of the growth kinetics decreases by ∼40%, while at low temperature there is more than two fold increase in the lag phase of the growth kinetics. Both the chemical environments evidently show to suboptimal growth rates (S1 Fig 7B), especially decrease in the amplitude in presence of glycerol suggests a stronger selection pressure.

Through DMS, DFEs obtained for all test environments show significant disparities as compared to reference environment i.e. 37°C (Fig 1C, S1 Table1). Position-wise distributions of fitness reveal differential enrichment and depletion along the length of the gene across all test environments (S1 Fig8). Notably, survival of mutants under elevated temperature condition and environments involving elevated temperature condition is greatly lowered (S1 Table1) indicating deleterious effects of elevated temperature condition. Among environment specific DFEs, compared to reference environment, at low temperature, survival of a subpopulation of mutants is compromised (Δn = −141) while average fitness (ΔF = 0.33) of survived mutants in the condition is increased. The increased average fitness is majorly due to compromised survival of the fraction of depleted mutants. As the natural host of GmR gene i.e. *S. marscens* requires lower growth temperature for optimal growth (27), the gene may have been evolutionary optimized to prefer lower temperatures thus exhibiting partial mutational robustness (n_pos_ = 40, n_neg_ = 9). Elevated temperature, imposes drastic deleterious effects on the survivabilities of mutants (Δn = −637) accompanied by increased average fitness of survived mutants (ΔF = 0.19). Lowered survivabilities are arguably due to protein misfolding (28) while because of prominent elimination of low fitness temperature sensitive mutants the average fitness of mutants is increased. The survival of high fitness mutants under elevated temperature condition may have resulted due to stabilization to a particular trait such as higher thermal stability.

Among chemical environments, both the chemical chaperones are found to exert mutational robustness as evident from retained survivabilities of mutants. In terms of fitness change, fitness gain is stronger in case of glycerol (ΔF = 0.42) than TMAO (ΔF = 0.21) (Fig1C). The mutational robustness supports the known mutational buffering as reported earlier (18). Notably, different sets of mutations are buffered by the two chemical environments (S2 Table) supporting known mechanistic difference in aiding protein folding (18,19). Further, at stringent Gm selection (25 μg/mL), compared to minimal Gm selection, TMAO treatment is found to exert mutational robustness (S1 Fig 9A). Remarkably, glycerol treatment exerts mutational robustness compared stringent (S1 Fig 9A) as well as minimal Gm selection (S1 Fig 9B). Collectively, among individual treatment of environments, low temperature and chemical chaperones induce mutational robustness while, elevated temperature exert deleterious fitness effects.

### Fitness effects of combinations of environments

Phenotypic effects of simultaneous action of multiple environments have been historically illusive to decipher (29). Using our experimental system, we set out to elucidate the effects of combinations of different sets of environments having prominent but contrasting effects i.e. elevated temperature and chemical chaperones. From growth kinetics of the model system, under combination of environments, it is apparent that TMAO is able to counteract the effects of elevated temperature while in case of treatment of glycerol, compared to growth kinetics at elevated temperature, the kinetics drastically slows down (S1 Fig 7C). Co-culture bulk competitions under simultaneous action of environmental conditions i.e. 42°C + TMAO and 42°C + glycerol reveal drastic decrease in survivability of mutants. Interestingly, average fitness of mutants is in case of 42°C + TMAO condition is higher than that for 42°C + glycerol condition. Contrasting effects of the combinations of environments are consistent with the low predictabilities fitness effects associate with simultaneous action of multiple environments.

In order to deduce relative influence of individual environments in the combinations, we quantitated their selection metrics against individual constituent environments. Compared against 42°C alone (Fig 2A), both 42°C + TMAO and 42°C + glycerol conditions show weaker differences (Fig 2A) than when compared against chemical environments (Fig 2B and C). This suggests that the effects of combinations of environments closely resemble to treatment of elevated temperature which has a strong influence in the combination. If considered as simple additive effects, for both the cases of combination of environments, the resultant difference in fitness do not seem to be a simple addition (Fig 2D and E). While addition of changes in fitness scores for the constituents environments weakly correlate with the fitness changes of combination of environments, among the two constituent chemical environments, glycerol treatment produces lower predictability in terms of the additive model (r=0.54) than TMAO (r=0.75). Collectively, while in terms of survivabilities of the mutants, elevated temperature has a strong influence, explaining the non-additive fitness effects would need mechanistic insights.

### Molecular constraints of underlying E2P effects

Fitness effects would also depend upon constraints of multiple aspects such as structural, folding and binding. Here, we used a set of molecular constraints describing multiple aspects of gene fitness (S3 Table). Apart from measure of perturbation of stability by mutations (ΔΔG), we use residue depth as an additional feature representing folding component. Core residues are known to play central role in protein stability (30); so mutations at the core of protein would be less tolerated (lower fitness) than the ones at the surface of the protein. Other structural features such as distances from dimer interface, residue flexibilities (B-factor obtained by PDB structure) describe conformational constraints of the protein. Differences between physico-chemical properties such as hydrophobicity (logP) and solvent accessibility of mutant and reference amino acids account for mutational perturbations of non-covalent interactions. Mutations near or at active site are more likely to have deleterious effects than mutations far from active site. Using this principle, mutational perturbations of ligand binding are accounted using a proxy of distance of residues of the protein from active site residue (D147) of the protein. Here, minimum distance between the atoms of the D147 and C-alpha atom of the residue is used to ensure maximum sensitivity. Conservation scores of the residues were included for accounting for culmination of structural, folding and binding constraints in the form of evolutionary rate per site.

Correlations of fitness scores with multiple molecular features quantitated in terms of Spearman’s rank correlation coefficients (ρ) reveal distinct classification of molecular constraints (Fig 3). Euclidian clustering along the axis of molecular features separates folding and binding constraints. Conservation scores, in particular, are best correlated with the fitness scores across different environments. Among negatively correlated molecular features, correlations with measures of protein stability estimates (ΔΔG) (31) and residue depth reveal that mutations with high stability perturbations or high residue depth show lower fitness and vice versa. Contrastingly, distances from sensitive active site and dimer interface of the protein are positively correlated with fitness scores. So mutations near active site or near dimer interface indeed show lower fitness. Other than top five molecular constraints, structural constraints such as residue flexibility, Δ(logP) per substitution and Δ(Solvent Accessible Surface Area) per substitution are mostly negatively but relatively weakly correlated to fitness scores across different environments. Overall, apart from conservation scores, folding and binding components stand out as the strongest structural predictors of the fitness effects across all environments; suggesting its major role in shaping the fitness landscape of GmR.

### Differential strengths of molecular determinants underlie environmental selection pressures

Alongside the classification of molecular constraints, Euclidean clustering (Fig 3) along the axis of environments reveals a distinct classification which in turn aligns with their effects on survivabilities i.e. Δn (Fig 3). Notably, correlation with conservation score is the highest in case of reference condition (37°C) than other environments; implying that the mutational tolerances in this environment closely resemble to that existed in evolutionary history of the gene. This supports our primary assumption to regard 37°C as a reference environment. Molecular features accounting for folding constraints i.e. ΔΔG and residue depth are best correlated in case of environments with conserved (least changed) survivabilities suggesting that the environments may induce allow survival of mutants with fitness changes that are aligned with the folding constraints of the protein. In other words, folding constraint is likely to be stronger in such environments than the environments with compromised survivabilities. Contrastingly, in terms of distance from active site and dimer interface, fitness scores of mutants from environments with lowered survivabilities are better correlated. This suggests that these factors particularly the best correlated between the two i.e. distance from active site is potentially a major fitness constraint.

Residue flexibility, interestingly, is negatively correlated with fitness scores of mutants from environments with lowered survivabilities and positively correlated with majority of rest of the environments. This implies that for environments that confer strongly deleterious fitness effects, survived mutants with higher fitness scores are more likely to have lower residue flexibility at mutation sites and vice versa. Elevated temperature conditions are known to promote thermostability by reducing the conformational plasticity of the proteins (32,33); thus explaining why residue flexibility emerges as a relatively strong constraint. Physico-chemical features of the substitutions i.e. Δ(logP) per substitution and Δ(Solvent Accessible Surface Area) per substitution, on the other hand, emerge as relatively weaker constraints; suggesting that perturbations of non-covalent interactions have relatively minor role in guiding fitness of mutants.

Although conservation score and molecular features describing folding and binding are seem to be relatively strong constraints of fitness effects, individually, they are weakly correlated (ρ<0.4) with the fitness scores. Assuming that the uncertainty in estimations of structural and predicted features is considerably low and considering low experimental noise in determination of fitness scores of GmR mutants, the weak correlations are consistent with known non-monotonic relationship between fitness scores with biophysical constraints (13). Additionally, the weak correlations can be a result of complex interplay among multiple factors such as well known couplings between folding and binding components (34,35) or with other interdependent factors (36). This exemplifies the underlying dependence of mutational tolerance on the inherent structural constraints of the protein. Considering the possible complex interplay of set of molecular constraints, we specifically focus on the strongest structural constraints i.e. folding and binding for further contextualizing the environmental effects.

### Dependence on folding and binding constraints

Protein folding and ligand binding are known to act as spandrels underlying fitness effects (34,35). Corroboratively, hereby, we demonstrated that the two factors potentially play central role in guiding the fitness of GmR mutants. Based on these molecular features, next, we made four subsets of mutants with distinct combinations of folding and binding states: proper folding and binding (FB), compromised folding and proper binding (cFB), proper folding and compromised binding (FcB) and compromised folding and binding (cFcB). Here, cF and cB indicates compromised folding (high ΔΔG) and compromised binding (low distance from active site) respectively; while F and B denote non-compromised folding (high ΔΔG) and non-compromised binding (high distance from active site) respectively. Median values of ΔΔG and distance from active site is used as threshold to bin the mutants in 4 equal sized subsets. For instance, subset cFcB represents mutants with ΔΔG values more than the median ΔΔG and distance from active site less than the median distances from active sites for all residues. Here, because the classification is based on values of molecular features inferred from crystal structure of GmR (37), to reduce the influence of the uncertainties involved, we exclude the mutations with values of molecular features falling between middle 20 percentiles from the classification. Notably, known important predictor of protein stability (30), residue depth is positively correlated with stability perturbation i.e. ΔΔG (ρ=0.44, S1 Fig 10). So mutants with high stability perturbation (cF) are highly likely to reside in the core regions of the protein while the ones low stability perturbation (F) are highly likely to lie on the surface of the protein.

Next, we calculated subset wise metrics of selection i.e. Δn and ΔF with respect to reference environment (37°C, 12.5 μg/mL Gm). Firstly, in terms of survivability i.e. Δn, except for treatments of chemical chaperones at 12.5 μg/mL Gm and treatment of glycerol at 25 μg/mL Gm, for rest all the environments, survivability of the mutants is drastically decreased (Fig 4A). Notably, for environments with decrease in levels of survivabilities, across subsets, Δn seem to show a pattern which is closely dependent on the binding and folding constraints. Survivability of the mutants seem to decrease in a following order FB > cFB > FcB > cFcB. The greater decrease in survivabilities with the compromise in folding and binding supports our primary basis for classification of mutants and indicates that fitness scores of mutants are closely dependent on folding and binding constraints.

In terms of change in average fitness (ΔF), with a clear exception of 37°C Gm 25μg/mL, for majority of environments average fitness of surviving mutants is greater than reference environment (Fig 4B). Stringent Gm selection in addition to decreasing survivabilities of mutants, seem to lower the fitness levels of mutants. Among subsets, surviving mutants from subset FcB in particular have substantially lowered fitness scores potentially due increased dependence on substrate binding and catalytic activity because of increased level of purifying selection. Combination of environments too show lowered average fitness across subsets; effect of which is particularly exemplified in FcB subset. For other environments, the average fitness of surviving mutants seem to be increasing in the order FB < cFB < FcB < cFcB. This suggests that the compromise of folding and binding allow exclusive survival of mutants with fitness advantage. While mutants at the core of the protein and near to active site (cFcB) have lowered survival, the survived mutants tend to possess relatively increased fitness advantage.

Elevated temperature conditions is an interesting example wherein the survivability of mutants is drastically decreased (Fig 4A) and the environmental section only allowed survival of mutants with fitness advantage (Fig 4B). Overall, FB subset is least affected while contrastingly cFcB is the most affected subset by across all environments. Mutational robustness by TMAO and glycerol is apparent by the retention survivability of mutants (Fig 4A) and increased fitness of the mutants (Fig 4B). In combination with elevated temperature, both survivability and average fitness of mutants is decreased. At stringent concentration of Gm, unlike TMAO treatment, glycerol treatment confers mutational robustness. Collectively, distinct effects based on underlying influence of folding and binding states reveal a unique pattern in survivabilities and relative changes in average fitness.

### Unique reshaping of environment specific fitness landscape

In order to better contextualize the contributions of coupling between folding and binding, we visualized the fitness landscape of GmR by lining it with the folding and binding components which evidently act as central constraints of mutational fitness (Fig 5A). So, regimes at the corners of the fitness landscape represent the subsets of mutants with different states of folding and binding i.e. FB, cFB, FcB and cFcB.

For reference environment, as expected from corresponding folding and binding states, we find that majority of high fitness mutations occupy FB regime whereas cFcB regime is majorly occupied by the deleterious mutants (Fig 5A). Notably, this pattern is conserved across all the test environments (Fig 5B-J). For reference environment, folding constraint produces a pronounced fitness cliff whereby mutants with above a threshold of ΔΔG (∼2 kcal/mol) are highly likely to show deleterious fitness. In case of selection at stringent Gm concentration, corroborating with dosage dependent effects, mutations close to the active site show a prominent decrease in fitness (Fig 5B). Here, the imposed higher load of Gm, seem to generate additional pronounced fitness cliff along the binding axis. The substantial change in the fitness landscape as a function of selection levels is consistent with known central role of purifying selection in molecular evolution (21).

Among physical environments, fitness landscape corresponding to low temperature condition show no peculiar regime wise distinction from that of reference environment (Fig 5C); capturing earlier noted weaker selection pressure (Fig 1C). Elevated temperature conditions - in accordance with imposed negative selection pressure - show selective survival of only mutants with enriched fitness associated with lowered survival of mutants from cFB and cFcB subsets (Fig 5D). Among chemical environments, mutational robustness imposed by TMAO and glycerol is evident in the form of close similarity with the fitness landscape of reference environment (Fig 5E,G). At stringent Gm concentration, chemical chaperone seem to increase the fitness of survived mutants lying at cFcB regime (Fig 5F,H) which are otherwise highly depleted (Fig 5B). For combinations of environments, in alignment with in previous analyses, fitness landscapes (Fig 5I,J) show substantially decreased survivabilities and contrasting average fitness levels i.e. particularly at cFcB regime glycerol treatment depletes the fitness of mutants more intensely than TMAO treatment. Overall, the distinct reshapings of the fitness landscapes under different environments reveal divergent fates and preferences for mutations of gene guided by differential strengths of key molecular constraints.

## Discussion

Large-scale elucidation of G2P interactions enabled by high throughput mutational scanning (38) has opened up new possibilities to comprehensively assess fundamental questions in molecular evolution. Here we investigate molecular level E2P interactions and their underlying mechanistic insights. Upon monitoring empirical fitness of a library of single site mutants of GmR, we characterized relative selection pressures imposed by sets of physical and chemical environments (Fig 1C). Central role of maintenance of cellular proteostasis is exemplified by the directionalities of the selection pressures which show a close dependence on the potential alterations of protein folding. This also supports the identification of stability perturbation and other protein folding related constraints as the one of the strongest constraints of fitness of GmR mutants.

Through our data, as in case of mutational scanning’s of other antibiotic resistant genes (21,22,39), we find that the changes in fitness landscapes of GmR are dependent on purifying selection level i.e. the dosage of antibiotic (Fig 1B). Corroborating with earlier studies (11,12), we demonstrate that mutational landscapes monitored under different environmental alterations of proteostasis are significantly different than that monitored under reference environmental conditions (Fig 1C). Among such environments, elevated temperature (42°C) exerts negative selection pressure underscoring known protein misfolding effects (28) and temperature sensitivity (17). Low temperature (30°C) conditions impose weak mutational robustness conforming to known non-deleterious effects on protein folding (40). Chemical chaperones, too, conforming with their known favorable effects on protein folding (18,19), exert mutational robustness. This shows that fitness effects in each environment on fitness landscape for GmR is dependent on kind of environmental alteration of cellular protein folding.

Remarkably, simultaneous action of environments, specifically environments with oppositely directed selection pressures i.e. elevated temperature and chemical chaperones are found to lead to lower predictable outcomes which underscore similar observation reported previously (11). In our study, estimation of selection pressures against constituent individual environments revealed that strongest among the constituent environments guide the overall selection pressure in the combination (Fig 2A). However in terms of average fitness, they have contrasting (Fig 1C) and non-additive effects (Fig 2D and E). The contrasting effects of chemical chaperones exemplified at the elevated temperature conditions may be a result of inherent mechanistic differences in thermodynamic assistance to the folding of proteins i.e. entropic assistance by TMAO and enthalpic assistance by glycerol (19). Additionally, effects of chemical chaperones seem to be a function of growth temperature i.e. at 37°C, supplementation of chemical chaperones lead to increase in average fitness of mutants while that at elevated temperature supplementation of chemical chaperones lead to decrease in average fitness of mutants. Such effects could be due to simultaneous selection for multiple traits which complicates the resultant selection pressure both qualitatively and quantitatively (29).

Next, in order to gain mechanistic insights into of the evident E2P interactions, we contextualized them in terms of molecular constraints of the fitness landscapes. Correlations of fitness scores with molecular constraints reveal their differential strengths for each environment (Fig 3). Conservation score - accounting for evolutionary rate per site - is best correlated with the fitness scores across all the test environments; thus substantiating the biological significance of the obtained empirical fitness values. Next to conservation score, stability perturbation (ΔΔG), and perturbation in binding (distance from active site) are best correlated with the fitness scores. Notably, however the strength of correlations are relatively weaker (ρ<0.4); suggesting possibility of non-monotonic relationships (13) and a complex interplay of interdependent constraints.

In our study, coupled factors, folding and binding constraints distinctly emerge as strong determinants of fitness effects; and through their spandrel like property (34). To understand the trade-off between the two, we visualized fitness of GmR mutants in a form of fitness landscape lined by folding (ΔΔG) and binding axes (distance from active site) (Fig 5A). Across all environments, corroborating with known trade-off in case of antibiotic resistant genes between folding and binding (35), combinations of folding and binding states seem to underlie resulting fitness effects. For instance, combination of weaker folding and binding constraints (FB) is associated with largely enriched fitness levels while stringent folding and binding constraints (cFcB) are associated with deleterious fitness levels of mutants (Fig 5A). In case of reference environment, folding constraint introduces a prominent limiting fitness cliff (at ΔΔG=∼2 kcal/mol). Whereas for test environments, although at variable degrees, common existence of limiting fitness cliffs along folding axis (Fig 5) underscores central role of protein folding and stability in molecular evolution (41). The universal dependence on folding component especially at minimal purifying selection also explains the conformity of evident fitness effects with known effects of environments on protein folding and proteostasis. Alongside, this study therefore reveals a unique possibility of controlling mutational fates based on environmental alteration of major constraints of fitness landscape. Notably, binding constraint imposes a limiting fitness cliff which is more pronounced at stringent concentration of Gm (25 μg/mL) (shown in Fig 5B) and from our data, it seems to be a feature dependent on level of purifying selection levels.

Collectively, from a simple experimental system consisting of a conditionally essential gene with evidently weaker constraints imposed by events prior to protein expression, we identify environment dependent differential fitness of mutations which are dependent on relative strengths of underlying molecular constraints. Major implication of this information lies in the improvement of our understanding of influence of environmental conditions on G2P interactions at molecular level. Further, environmentally induced variability can potentially contribute a requisite nuance in predictive models used for challenging task of inferring phenotypic outcomes from genomic variants(15,16); making them more robust. In future, such study of comprehensive environment specific fitness landscapes can be potentially extended to multiple mutations to monitor combined effects of epistasis - arguably the major predictive factor in molecular evolution (42), - more variety of environments as well as assessing fitness landscapes of multiple proteins in gene regulatory networks.

## Materials and Methods

### Minimal inhibitory concentration (MIC) assays

Primary culture was prepared by inoculating (1% v/v) *E. coli* (K-12) in culture media (Luria-Bertani (LB) broth (HiMedia) containing 100μg/mL, ampicilin (Sigma) and 0.1% Arabinose (Sigma)) and incubating at 37°C for 18 hrs. The primary culture was inoculated at OD_600_ of 0.025 in culture media containing a range of Gm (Sigma) concentrations from 6.25 to 400 μg/mL with 2 fold increase at each increment (in 96-well storage plates). The assay plates were incubated at 37°C for 18 hrs before measuring growth (OD_600_) in Tecan microwell plate reader.

### Growth assays

*E. coli* (K-12) harboring pBAD-GmR is grown in culture media (LB media containing 100 μg/mL and ampicilin 0.1% Arabinose) for ∼18 hr. Primary culture was used as an inoculum (∼0.01 OD) for the growth assays. Growth assays in different environments were carried out using Bioscreen C kinetic growth reader. The growth parameters were obtained by fitting absorbance data to five parameter Logistic equation.

### Co-culture bulk competition assay

SSM library of GmR was constructed by PCR based site directed mutagenesis using primers with degenerate codons (NNK). For detailed information regarding the mutagenesis, please refer to Supporting methods described in Bandyopadhyay et al. (18). For co-culture bulk competition assays, the mutation library cloned in pBAD vector was transformed into *E. coli* (K-12). Primary culture was prepared by inoculating pool of SSM library (1% v/v) in culture media (LB media containing 100 μg/mL ampicilin and 0.1% Arabinose) at 37°C for 18 hrs. A competition was carried out at the secondary culture where primary culture in inoculated at OD_600_ of 0.025 and incubated for 18 hrs. Physical environmental conditions were created by carrying out the bulk competitions at 30°C (low temperature) or 42°C (elevated temperature). Chemical environmental conditions were created by supplementing either TMAO (250mM) or glycerol (250mM) in the culture media of competition assay. Biological replicates were made by carrying out independent co-culture bulk competitions of the mutant libraries. For measuring fitness of mutants in a particular environmental condition, bulk competition under Gm selection (selected pool) (as shown in Fig 1A) was carried out. An independent bulk competition was carried out at 37°C in the absence of Gm (unselected pool) which serves as a reference for calculating preferential enrichments.

### Deep sequencing

At the end of bulk competition assays, cells are pelleted and plasmid is purified. Amplicons were generated by a short PCR (initial denaturation: 95°C for 3 min, denaturation: 95°C for 1 min, annealing: 60°C for 15 sec, extension: 72°C for 1 min, final extension: 72°C for 10 min) using high fidelity KAPA HiFi DNA polymerase (cat. no. KK2601). High template concentration (1 ng/μl) and 20 cycles were used to reduce potential PCR bias. Multiplexing was carried out using flanking barcoded primers (4 forward, 4 reverse, sequences in S1 Table 2). Amplicons of barcoded samples were grouped in equimolar concentration and gel purified. A dual index library for each such set was prepared using Truseq PCR-free DNA HT kit (Illumina Inc. Cat no. F-121-3003) and sequenced using paired end (300 X 2) chemistry on Illumina Miseq platform. Raw sequencing data is available upon request.

### Estimation of fitness scores from deep sequencing data

Analysis of sequencing data was carried out by using *dms2dfe* (43) - a comprehensive analysis pipeline exclusively designed for analysis of deep mutational scanning data. Through *dms2dfe* workflow, output files from the sequencer (.fastq) were demultiplexed using *ana0_fastq2dplx* module of *dms2dfe*. Average read depth of each demultiplexed sample was ∼1X10^5^. Next, though *dms2dfe*’s modules namely *ana0_fastq2sbam*, sequence alignment was carried out using Bowtie2 (44) followed by variant calling through *ana1_sam2mutmat* module which utilizes *pysam* libraries (45). A variant is called only if average Q-score of the read and that of the mutated codon is more than 30. Additionally a cut off of 3 reads per variants is used to filter out anomalous low counts. As a result a codon level mutation matrix of counts of mutations is generated. Codon level mutation matrix is then translated to amino acid level (based on the codon usage bias of the *E. coli*). For each experimental condition, counts of ∼2000 individual mutants were quantified (S2 Table). Raw sequencing data is available at Sequence Read Archive (SRA) as a BioProject: PRJNA384918.

Through *ana2_mutmat2fit* module of *dms2dfe*, counts of mutants are first normalized by the depth of sequencing at each position of the gene. Then preferential enrichments which are log (base 2) fold change of counts of the mutants in pool selected in presence of Gm against unselected (0 μg/mL Gm) reference pool are estimated. Here, preferential enrichment of a mutant serves as a proxy for its relative fitness and hence we simply refer it as ‘fitness’. Alongside significance levels of the preferential enrichments are obtained by Wald test through DeSeq2 workflow (46) (S2 Table).

### Classification of mutants based on comparison of DFEs

In order to assess the difference in fitness scores in a given comparison of DFEs of two environments, we assigned a statistical threshold to reduce the influence of experimental and biological noise. The mutants with higher fitness scores in test environments than control environment are classified as ‘positive’ and while mutants with lower fitness scores in test environments than control environment are classified as ‘negative’. We use replicates of test and control environments to get thresholds of difference in fitness scores such that the differences in fitness scores between the environments (‘signal’) is greater than the differences within the replicates (‘noise’). From difference between fitness scores of replicates, two distributions corresponding to replicates of test and control environment are obtained. μ_test_ and μ_control_ are means and σ_test_ and σ_control_ are standard deviations of such distributions for test and control environments respectively. So, the thresholds to segment the mutations into ‘positive’ or ‘negative’ is determined to be equal to 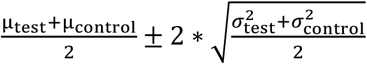. Number of mutants undergoing positive effects is denoted as n_pos,_ while number of mutants with negative effects is denoted as n_neg_.

### Molecular features of GmR

Mutant stability perturbations (ΔΔG) are predicted by PoPMusic (31) server. Conservation score is acquired from ConSurf (7) server. *MSMS* libraries (47) were used for calculations of residue depth from surface of protein. Distances between atoms of GmR are measured using various modules of Biopython package (48). Physico-chemical properties of the amino acids such as logP and pI were retrieved from PubChem (49) and ChemAxon (http://www.chemaxon.com). Molecular features of mutations used in the study are included in S3 Table.

## Supporting information

**S1 Text. Supporting Figures and Tables.**

**S2 Table. Fitness scores of mutations in different environments.**

**S3 Table. Molecular features of mutations.**

## Abbreviations

G2P: Genotype to Phenotype
E2P: Environment to Phenotype
Gm: Gentamicin;
GmR: Gentamicin Resistant gene;
DMS: Deep Mutational Scanning;
DFE: Distribution of Fitness Effects;

## Authors’ contributions

KC designed the study. RD carried out DMS including bulk competitions, DNA purification and sequencing. RD and KC carried out the analysis of sequencing data. RD and KC wrote the manuscript. R.P and G.J. helped in the amplicon sequencing. All authors read and approved the final manuscript.

## Acknowledgments

We acknowledge CSIR for its funding through BSC0124 project and infrastructural support from CSIR IGIB. R.D. acknowledges UGC for graduate funding. We thank members of KC lab for critically reviewing the manuscript.

